# Decoupling of GABA and Glutamine-Glutamate Dynamics and their role in tactile perception: An fMRS Study

**DOI:** 10.1101/2024.11.28.625809

**Authors:** Duanghathai Pasanta, Helen Powell, Nauman Hafeez, David J Lythgoe, Nicolaas A Puts

**Affiliations:** Department of Forensic and Neurodevelopmental Sciences, Institute of Psychiatry, Psychology, and Neuroscience, King’s College London, London, London SE5 8AB, United Kingdom; Department of Radiologic Technology, Faculty of Associated Medical Sciences, Chiang Mai University, Chiang Mai 50200, Thailand; Department of Neuroimaging, Institute of Psychiatry, Psychology, and Neuroscience, King’s College London, London, London SE5 8AB, United Kingdom; MRC Centre for Neurodevelopmental Disorders, New Hunt’s House, Guy’s Campus, King’s College London, London, SE1 1UL London, United Kingdom

**Keywords:** fMRS, tactile processing, GABA, glutamate

## Abstract

Tactile processing is fundamental for our daily lives. In particular, adaptation, the mechanism by which neural (and behavioural) responses change due to repeated stimulation, is key in adjusting our responses to the environment and is often affected in neurodevelopmental conditions such as autism and ADHD. While GABA and glutamate—the main inhibitory and excitatory neurotransmitters— are known to be fundamental for encoding sensory input, we know little regarding the dynamic responses of the GABA and glutamatergic systems during tactile processing. Here, we examine how GABA and glutamine+glutamate (Glx) *in vivo* dynamics change during repetitive tactile stimulation and how these changes relate to tactile perception in a healthy population, using functional magnetic resonance spectroscopy (fMRS). Our study showed that repetitive tactile stimulation induced a decoupling between GABA and Glx during the first stimulation blocks as suggested by a negative correlation between GABA and Glx, which changed from a positive correlation at baseline. Subsequently, a multivariate time series analysis showed a predictive temporal relationship between Glx and GABA, showing that changes in these metabolites are temporally linked with an estimated lag of 6 seconds informing on a complex metabolite response function. The absence of gross metabolite change suggests that Glx and GABA adjust in relation to each other in response to repeated tactile stimulation. Furthermore, individual differences in the changed GABA and Glx levels correlated with perceptual measures of touch. Together, our study highlights the complex relationship between GABA and glutamate in tactile processing and demonstrates that experience-dependence plasticity induces a decoupling between these key metabolites. Further study into their dynamic interplay may be key to understanding adaptation as meso-levels in the brain and how these mechanisms differ in neurodevelopmental and neurological conditions.

## Introduction

Touch plays an important role in our everyday lives. Beyond sensing and interacting with the world around us, touch is also crucial for fostering interpersonal relationships and is ultimately associated with our emotional well-being.^1,2^ Atypical tactile processing is also common in neurodevelopmental and affective disorders. For example, differences in touch perception have been associated with core symptoms of neurodevelopmental conditions such as autism and attention deficit hyperactivity disorder (ADHD).^3–5^ Given the importance of touch, and the impact of atypical touch, understanding how touch is processed in the brain is of high relevance.

Glutamate and GABA are the main excitatory and inhibitory neurotransmitters in the brain, respectively. It is well established that glutamate and GABA are key modulators of sensory processing.^6^ Through a delicate balance between excitatory (E; predominantly glutamatergic) and inhibitory (I; predominantly GABAergic) function, GABA and glutamate facilitate and tune neuronal responses for optimal information encoding and subsequently generate context-appropriate responses to incoming sensory information.^6–9^ For example, the GABA and glutamate systems play key roles in the encoding of tactile frequencies^10,11^, intensity^12–14^, and spatial information.^15–17^

Beyond the immediate encoding of tactile information, we respond to changes in our sensory environment through adaptation and habituation.^18^ Changes in synaptic strength and plasticity are crucial for optimising sensory responses and adapting to stimuli, for example, through synaptic strengthening or pruning to enhance acuity and precision in response to repeated stimuli.^19,20^ It has been robustly demonstrated that passive exposure to repetitive tactile stimulation over a few minutes leads to synaptic plasticity at the cellular level and corresponding perceptual improvement via modulation of E/I.^21–24^ Repetitive tactile stimulation has also been shown to reduce cortical response amplitudes in animal models,^25,26^ and enhance perceptual sensitivity in humans.^24,27–29^ An animal study using a GABA antagonist demonstrated that inhibiting GABAergic lateral inhibition increased receptive field size, thereby worsening tactile discrimination.^30^ However, how the GABA and glutamate systems respond during sensory stimulation remain uncertain, and have not yet been examined in humans.

Magnetic Resonance Spectroscopy (MRS) is currently the only technique that can be used to measure markers of GABA and glutamate *in vivo*. Prior studies measured GABA and glutamate in tactile processing relatively statically, rather than investigating the dynamic response of the E/I system. Even studies manipulating the GABA or glutamate system through pharmacology, or repeated stimulation, have determined their effects in a pre/post design.^31,32^ Beyond investigating changes in GABA or glutamate in response to repeated stimulation, it is well-known that the GABA and glutamate systems interact both neuronally and metabolically.^33^ However, it is not known to what extent their dynamics impact one another throughout sensory stimulation, or whether the dynamic response to sensory stimulation is reflective of perceptual capacity. While some studies have correlated static levels of GABA and glutamate to perceptual thresholds, the capacity of the system to change is likely more reflective of behavioural outcomes. This gap in knowledge highlights the need for more research to establish whether the dynamic nature of GABA and glutamate responses to touch correlate with or predict tactile sensitivity in behavioural assessments. This is particularly relevant to those neurodevelopmental conditions where both E/I and sensory processing, appear to be affected.^34–36^ Understanding these associations could provide deeper insights into the neurobiological mechanisms underlying sensory processing and potential GABA or glutamatergic intervention.

Functional magnetic resonance spectroscopy (fMRS) is an emerging non-invasive technique that adds functional aspects to ^1^H-MRS by assessing neurometabolite levels while participants perform a task or are exposed to a stimulus. Since the fMRS signal is not cell-compartment specific, MRS-measured GABA and glutamate are considered as overall, and indirect indicators of inhibitory and excitatory activity within a brain region. While very few studies have investigated the tactile domain using fMRS, a recent animal study demonstrated that prolonged whisker stimulation induced MRS-measured GABA and glutamate changes, reflecting neuronal activities of excitatory and inhibitory processes as assessed by two-photon imaging in awake mice.^37^ One study in humans showed that high-frequency stimulation-induced perceptual learning and reduced GABA levels.^10^ While previous studies have examined GABA and glutamate separately, there remains a limited understanding of their interrelationship. Specifically, it is unclear how these neurotransmitters influence each other and what factors influence their response.

Thus, fMRS is uniquely positioned for studying dynamic changes in glutamate and GABA in response to tactile stimulation. In this study, we used passive repetitive sensory stimulation to establish the link between tactile processing and glutamate/GABA dynamics in the somatosensory cortex, as this has previously been shown to induce plastic changes in the cortex using fMRS.^10,38^ To examine their impact, we then also related changes in GABA and glutamate levels to tactile perception. Here, we hypothesised that following repetitive sensory stimulation, there would be a reduction in GABA levels to facilitate perceptual plasticity and an increase in glutamate due to energetic response.

## Materials and methods

### Participants

A total of 20 participants participated in the study (mean age (SD) = 25.75 (5.79) years old; 12 females). Written informed consent was obtained from all participants under the approval of the local institutional ethics committee at King’s College London. Participants had no current neurological or physiological diagnoses as per oral confirmation. Handedness was verified with the Edinburgh Handedness Inventory.^39^ The Glasgow Sensory Questionnaire was used to assess sensory sensitivities, where a higher score is indicative of greater sensory difficulties and has previously been shown to be associated with higher autistic traits.^40^ We investigated autistic traits using the Autism Spectrum Quotient 10.^41^

### Vibrotactile measures

A battery of vibrotactile tasks was used to assess the perceptual sensitivity of each participant with a total duration of 35 minutes. Participants were asked to sit comfortably and position the index and middle fingers of their left hands on a two-digit vibrotactile stimulator shaped like a computer mouse (Cortical Metrics, North Carolina, USA). Participants responded using a computer mouse with their contralateral hand. All tasks have been described previously.^42^ Stimuli were delivered within the flutter range (frequency = 0–50 Hz, amplitude = 0–350 μm), controlled by the BrainGauge program on a laptop computer. At the beginning of each task, a short explanation and three practice trials were provided to confirm that participants understood the task.

A total of eight vibrotactile tasks were assessed for all participants, namely: simple reaction times, choice reaction times, static detection thresholds, dynamic detection thresholds, amplitude discrimination without adaptation and with single-site adaptation, simultaneous frequency discrimination, and temporal order judgment (Supplementary Fig.1). For all but the reaction time tasks, thresholds were determined using a staircase tracking paradigm as previously reported.^42^ The vibrotactile measurement data were exported and then analysed using a custom R package (available at: https://github.com/HeJasonL/BATD). The difference between dynamic and static detection thresholds was calculated to assess a proxy measure of feedforward inhibition, and the difference between amplitude discrimination thresholds with and without adaption was also calculated to assess the effects of sensory adaptation.^42^

### fMRS methods

All MRI data were acquired on GE 3 T Signa Premier (GE Healthcare, Chicago, Illinois) with a 48-channel head coil. T1-weighted structural images (MP-RAGE; 1 mm^3^, TE/TR = 2.96/2658.96 ms, TI = 860 ms, flip angle = 8°) were acquired from each participant for voxel placement. Due to the relatively low concentrations of GABA and glutamate within the brain, we used the J-difference editing MEscher–GArwood Point RESolved Spectroscopy (MEGA-PRESS) sequence for quantification of GABA and Glx (glutamine + glutamate, which cannot be separated at 3 T).^43–45^ For MEGA-PRESS, the GABA signal at 3 ppm is co-edited with the macromolecule signal and will be referred to as GABA+ from this point onward. The acquisition parameters for MEGA-PRESS were TE/TR = 68/2000 ms, voxel size = 3x3x3 cm^3^, data points = 2048, Spectral width = 2000 Hz. Spectral editing pulses with bandwidths of 60 Hz were applied at 1.9 ppm for edit-ON and 7.5 ppm for edit-OFF frequencies, symmetrical to the water resonance at 4.7 ppm. A chemically selective saturation (CHESS) water suppression pulses were used for water suppression. A two-step phase cycle was used for the excitation pulse, and individual transients were saved for later fMRS data processing.

Two MEGA-PRESS scans were acquired consecutively from the same voxel placed in the right somatosensory region. Each MEGA-PRESS acquisition was divided into resting-state fMRS (no stimulus presentation) for 180 transients (REST; 6 mins) followed by passive tactile stimulation for 180 transients (FUNC; 6 mins), leading to a total of 360 transients. The MEGA-PRESS acquisitions were repeated to mitigate the impact of scanner drift and motion.

Tactile stimulation was delivered using a 3D-printed MRI-compatible tactile device (**Error! Reference source not found.**D), where contact with the skin was made using air-pressure-driven plastic probes to the participants’ index and middle fingers simultaneously. Each probe was driven by an air pressure of 2 Bar. During the experiment, participants received passive tactile stimulation without requiring a behavioural response. Participants were instructed to focus on the sensation of the stimulation. Throughout the FUNC period, participants received 2 seconds of 3 Hz tactile co-stimulation on the glabrous skin of the left-hand index and middle fingers, interleaved with 2 seconds of no stimulus presentation.

### fMRS analysis

All spectral analyses were carried out using a custom version of Gannet (version 3.3.2) using default post-processing settings, with Robust Spectral Registration for phase and frequency alignment.^46,47^ Voxel co-registration and tissue segmentation based on the structural T1-weighted images were done to determine the fractional tissue composition using SPM12.^48,49^ Voxel consistency across all participants was obtained with a function available in Osprey.^50^ GABA and Glx concentrations were quantified in institutional units (i.u.) in ratio to unsuppressed water signal within the same voxel and corrected for tissue fraction, to account for intrinsic metabolite differences in grey matter and white matter.^51^ Spectra were analysed in two ways: block and sliding window analyses. A schematic representation of the fMRS analyses is shown in Fig. 1. Data quality is reported in line with MRSinMRS in Supplementary Table 1.^52^

**Figure 1.**
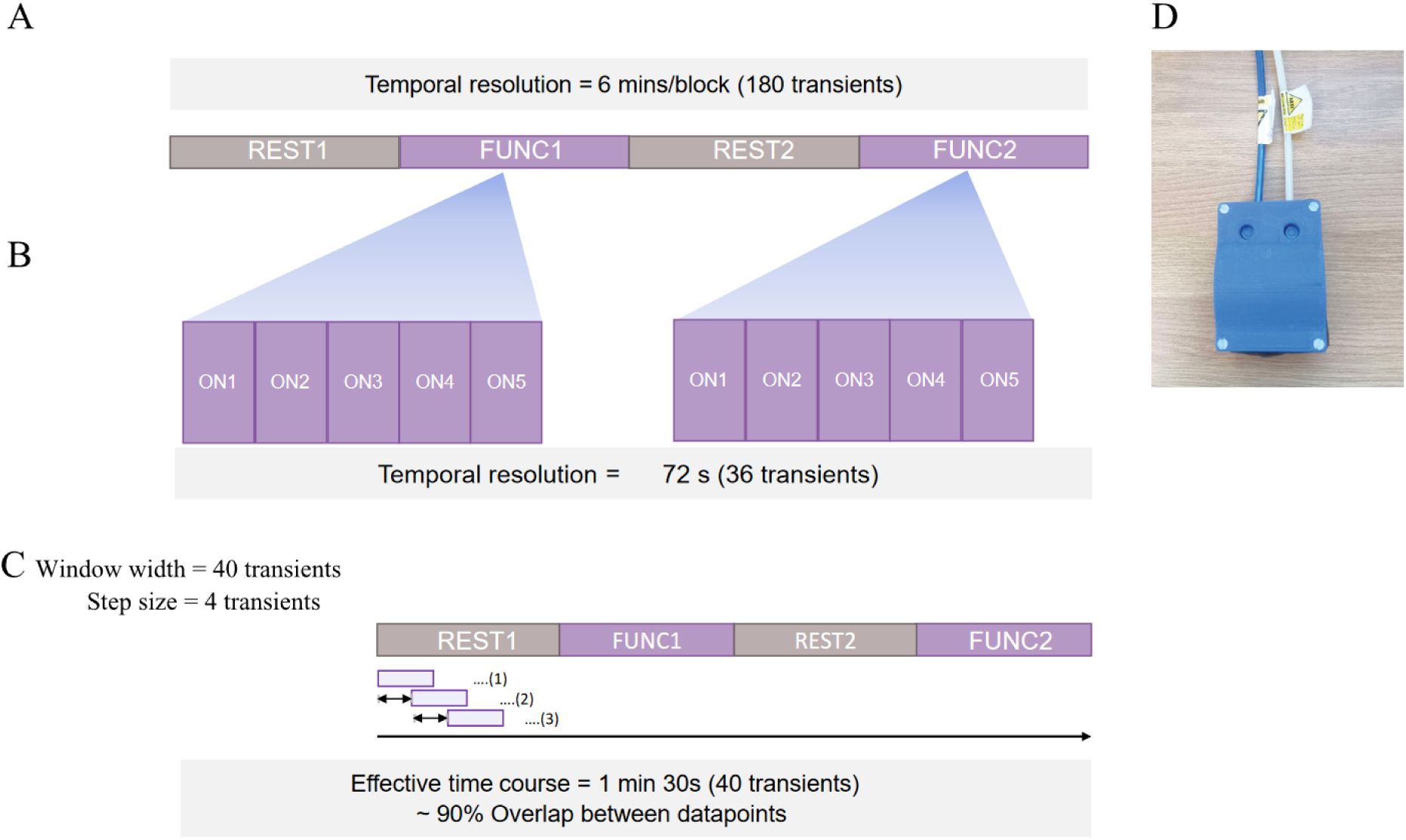
Schematic representation of fMRS analysis approaches in the current study. (A) Block analysis, (B) Subblock analysis where each FUNC block was divided into five subblocks (ON1–ON5) to gain higher temporal resolution, (C) Sliding window analysis of window width of 40 transients and step size of 4 transients, (D) 3D-printed MR-compatible vibrotactile stimulator device.

MRS data quality assessment was conducted by visual inspection for severe artefacts, and data points were excluded if the FWHM of NAA was > 20 Hz or the % fit error of that metabolite was more than 65%. Extreme outliers were then identified and removed using a boxplot method, where values above the third quartile (Q3) + 3×interquartile range (IQR) or below the first quartile (Q1) - 3×IQR were considered extreme outliers.

### Block analysis

The data were analysed in blocks based on how stimuli were presented, with 180 transients for each block (temporal resolution: 6 minutes) (Fig. 1A). We refer to the rest block and functional block of the first MEGA-PRESS acquisition as REST1 and FUNC1, and the rest block and functional block from the second MEGA-PRESS acquisition as REST2 and FUNC2. We also divided each FUNC block into 5 sub-blocks of 36 transients each to achieve higher temporal resolution (Fig. 1B). These sub-blocks were labelled as ON1–ON5 for each FUNC block, with a temporal resolution of 1 min 12 s.

### Sliding-window analysis

As an exploratory technique, and due to the likely metabolite response lag, spectra were also analysed using sliding-window analysis to gain a higher number of effective time points and to assess how metabolite levels change over time (Fig. 1C). This was done by averaging the transients over a window width of 40 transients as we previously established.^53^, then shifting this window forward with a 4 transients step size, repeated until the end of each MEGA-PRESS acquisition. This allows for an effective temporal resolution of 1 min 10 s (40 transients) and an effective time course of 8 s (4 transients).

### Statistical analysis

All statistical analyses were performed in R version 4.2.3.^54^ The normality of data was assessed by a Shapiro-Wilks test which in most cases reported a non-normal distribution. As a result, we used non-parametric and bootstrap approaches for analyses.

### Metabolite differences

The differences between metabolite levels obtained with block analysis and subblock analysis were assessed using a Kruskal-Wallis test, with a pair-wise post-hoc test using the Wilcoxon signed-rank test to obtain p-values, which were corrected for multiple comparisons using the FDR correction. Corrected p-values of < 0.05 were considered statistically significant.

For sliding window analysis, Bayesian analyses were performed to assess the effect of block using *brms*^55^ and *bayesanova.*^56^ To account for autocorrelation between data points obtained for each window, an additional autoregressive (AR) term was used. The Bayesian formula was as follows:

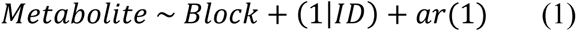

Weakly informative priors were applied across all Bayesian analyses.

Then, the Range of Practical Equivalence (ROPE) test was used to determine whether the observed effect could be considered practically significant.^57,58^ The ROPE test involved testing whether the 89% Highest Density Interval (HDI) of the posterior distributions fell within the range of negligible effect size or ROPE region; where ROPE was −0.1 to 0.1 times the standard deviation of the outcome variable.^58^ The 89% HDI was chosen based on its recommendation as a better choice than the 95% HDI.^59,60^ For the ROPE test, if the 89% HDI fell entirely outside the ROPE, the effect was considered practically significant. Conversely, if the 89% HDI fell fully within the ROPE, the effect was considered practically equivalent to null, implying the effect or difference was negligible. If the 89% HDI fell only partly within the ROPE, the result was inconclusive, indicating that more data might be needed to form a definitive decision.

Beyond looking at individual metabolites, we then examined whether interdependencies between GABA+ and Glx changed over time. Vector Autoregressive Model (VAR) analyses were carried out for sliding window data using the *vars* package.^61,62^ GABA+ and Glx were averaged across each time point to create time series data using the *tseries* package.^63^ Based on the tests of normality, the mean value was used for GABA+ since the data were normally distributed, and the median was used for Glx since it was not normally distributed. The optimal lag order for each model was determined using the *VARselect* function.^61,62^

To investigate whether GABA+ and Glx changes affected each other over time, the models were then tested for interdependencies between variables within the models (GABA+ and Glx) using a Granger-causality test.^64,65^ For example, if a time series variable A Granger-caused time series variable B, it indicates that the inclusion of time series variable A provides significant information for the future prediction of time series variable B.^66^ We then plotted the Impulse Response Functions (IRF) of each VAR model based on our data to investigate the temporal evolution of GABA+ and Glx to an impulse change in either GABA+ or Glx. The IRF represents the effect of an impulse (or a shock) to one variable on other variables in the same VAR model over time.^66^ For example, it can illustrate how an impulse change for GABA+ influences the evolution of Glx. For IRF plots, default confidence intervals of 95% were used, with a maximum of 20 lags, where each lag represented one transient (2 s), allowing us to achieve a 2 s effective resolution.

### Correlation analysis

To investigate the dependence of changes in neurometabolites and individual differences in metabolite levels at baseline, we employed Spearman correlation for non-normally distributed data (ρ: Spearman correlation coefficient) and Pearson correlation for normally distributed data (R: Pearson correlation coefficient), as determined by the test of normality. The variables of interest included the correlations between GABA+ and Glx for all fMRS analysis approaches, to evaluate how GABA+ and Glx changes in relations to each other under different conditions. To assess how individual baseline metabolite levels influence the magnitude of metabolite changes, we examined the correlations between baseline metabolite levels (REST1) and percentage change (%change) for each fMRS condition. The %changes were calculated by dividing the metabolite changes in each condition by the baseline condition (REST1) and multiplying by 100. Finally, we assessed the correlations between metabolite changes and vibrotactile measures to assess whether changes in GABA+ and Glx were related to measurable behavioural outcomes. Spearman or Pearson correlation analyses were used based on tests of normality here as well. Due to the exploratory nature of these measures, we did not correct for multiple comparisons in these analyses.

## Results

Descriptive statistics of participants’ demographics and questionnaire scores are shown in Supplementary Table 2. Averaged MRS spectral quality metrics after exclusions are shown in Supplementary Table 3. Averaged fMRS spectra and voxel positioning consistency across all participants are illustrated in Supplementary Fig. 2.

### No significant metabolite differences across conditions

Based on visual inspection, there were slight increases in both GABA+ (i.u.) and Glx (i.u.) during FUNCs compared to REST1 (Fig. 2A). However, the Kruskal-Wallis test did not show any statistically significant differences. Similarly, subblock analysis revealed no significant differences in both GABA+ (i.u.) and Glx (i.u.) under any condition (Fig. 2B). No noticeable patterns were observed within participants. The same lack of significant difference was observed for metabolites in ratio to creatine, suggesting no influence from the choice of reference (Supplementary Fig. 3).

**Figure 2.**
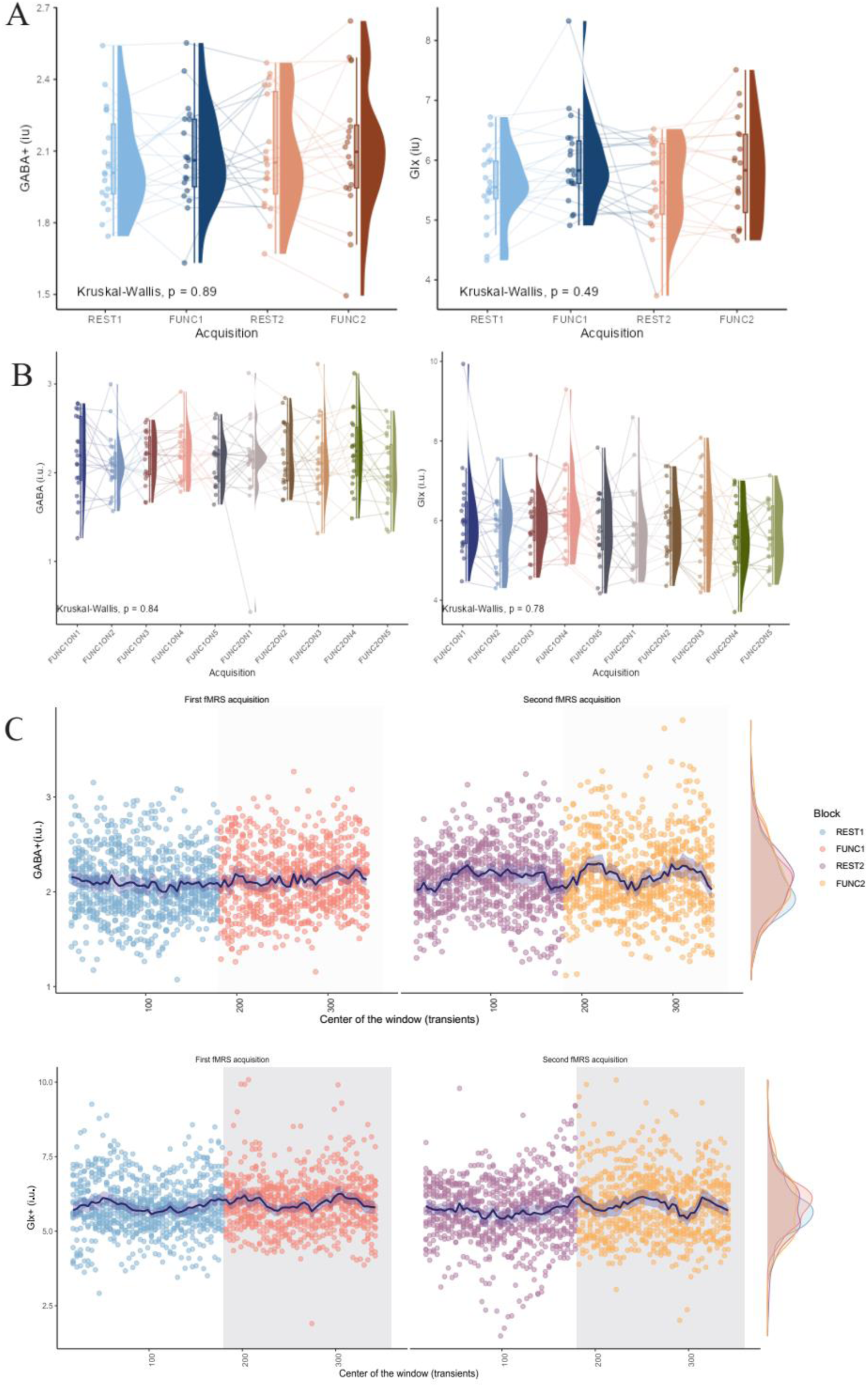
Raincloud plot of (A) block and (B) subblock analysis results. The grey lines in (A) and (B) connect data points within the same participant.

Based on visual inspection, the sliding window analysis showed no noticeable patterns across conditions (Fig. 2C**Error****! Reference source not found.**). The ROPE analyses from the Bayesian linear mixed model also agree, with the posterior distribution falling within the ROPE range of 60-85% (Fig. 3) and that most block effects are small with a low percentage of significance (Supplementary Table 4). These results suggest that the effect of blocks is considered to have an undecided significance for both GABA+ (i.u.) and Glx (i.u.).

**Figure 3.**
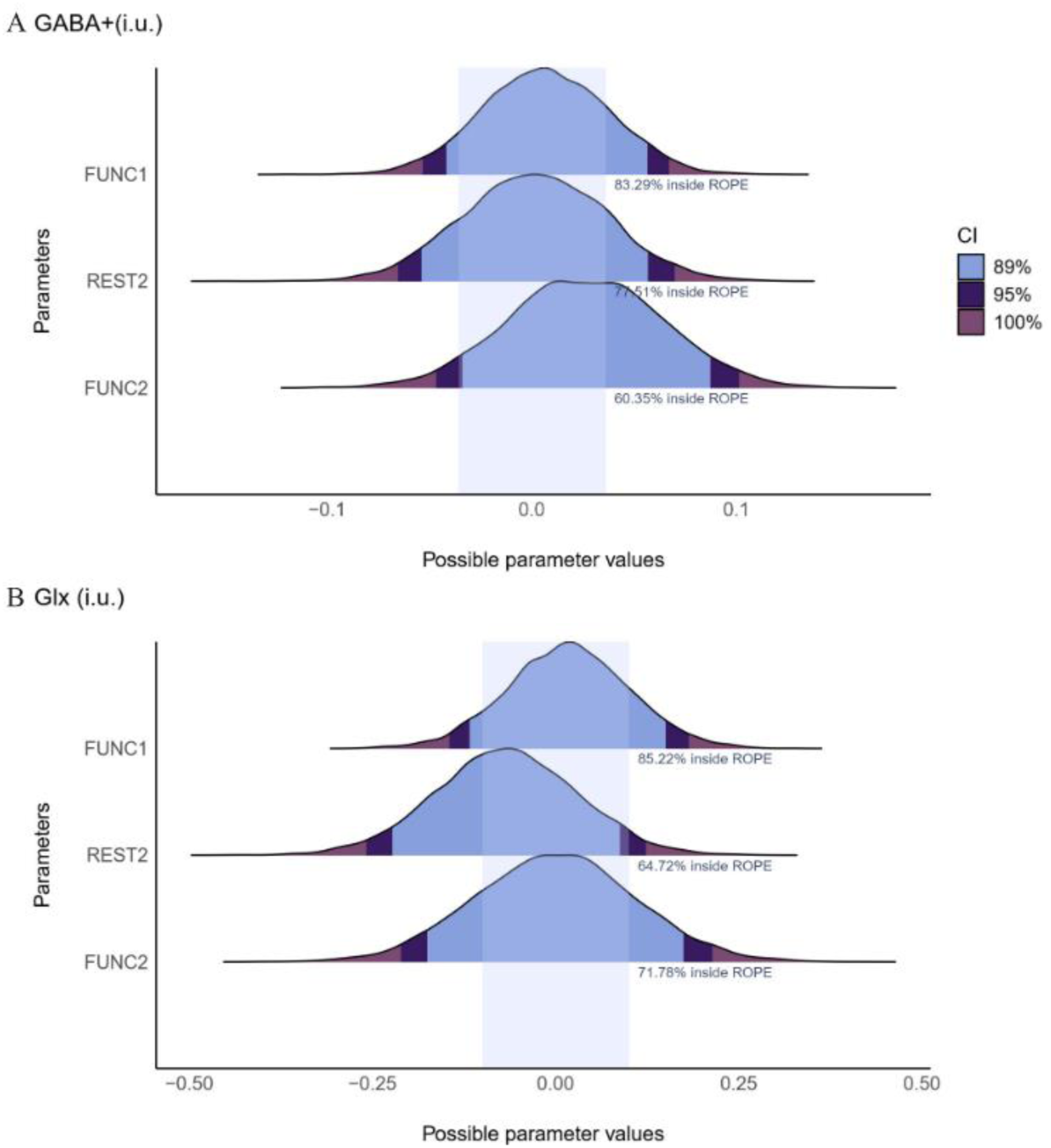
Posterior distribution plots of metabolite levels at each block with different shades represent the range within the posterior distribution that contains a specified proportion (89%, 95% and 100%) of the probability density. The superimposed blue boxes show the ROPE region (−0.1 to 0.1 times the standard deviation of the outcome variable which denotes the range considered practically equivalent to a negligible effect.^67^ The percentage displayed shows the portion of the 89% HDI of the posterior distributions that fall within the ROPE.

### Glx predicts GABA+ time series for FUNC1

Results from the VAR analysis of data obtained through sliding window analysis showed that the optimal lag order is p = 1 for all models. The Granger-causality between GABA+ (i.u.) and Glx (i.u.) was tested for each VAR model. The results indicated that for the VAR model including only data during FUNC1, the prediction of the GABA+ (i.u.) time series significantly improved with the inclusion of the Glx (i.u.) time series (F = 4.152, p = 0.045). Full results can be found in Supplementary Table 5.

### IRF plot suggested Glx increase after GABA+ impulse during FUNC1

IRF plots for VAR models of all data and only FUNC1 data are shown Fig. 4, while the remaining IRF plots for other data blocks are presented in Supplementary Fig 6. An impulse represents a change of one standard deviation of GABA+ (i.u.) or Glx (i.u.), where the response can be considered significant when the error band does not include the zero value. Overall, the IRF plots indicate the same trend of an increase in Glx following a GABA+ impulse and a decrease in GABA+ (i.u.) levels following a Glx (i.u.) impulse. Notably, the VAR model of FUNC1 data shows a significant increase in Glx (i.u.) levels for 3 lags (6s) after a GABA+ (i.u.) impulse, which then gradually returns to baseline, as depicted in Fig. 4B. Full results of all IRF plots can be found in Supplementary Fig.4.

**Figure 4.**
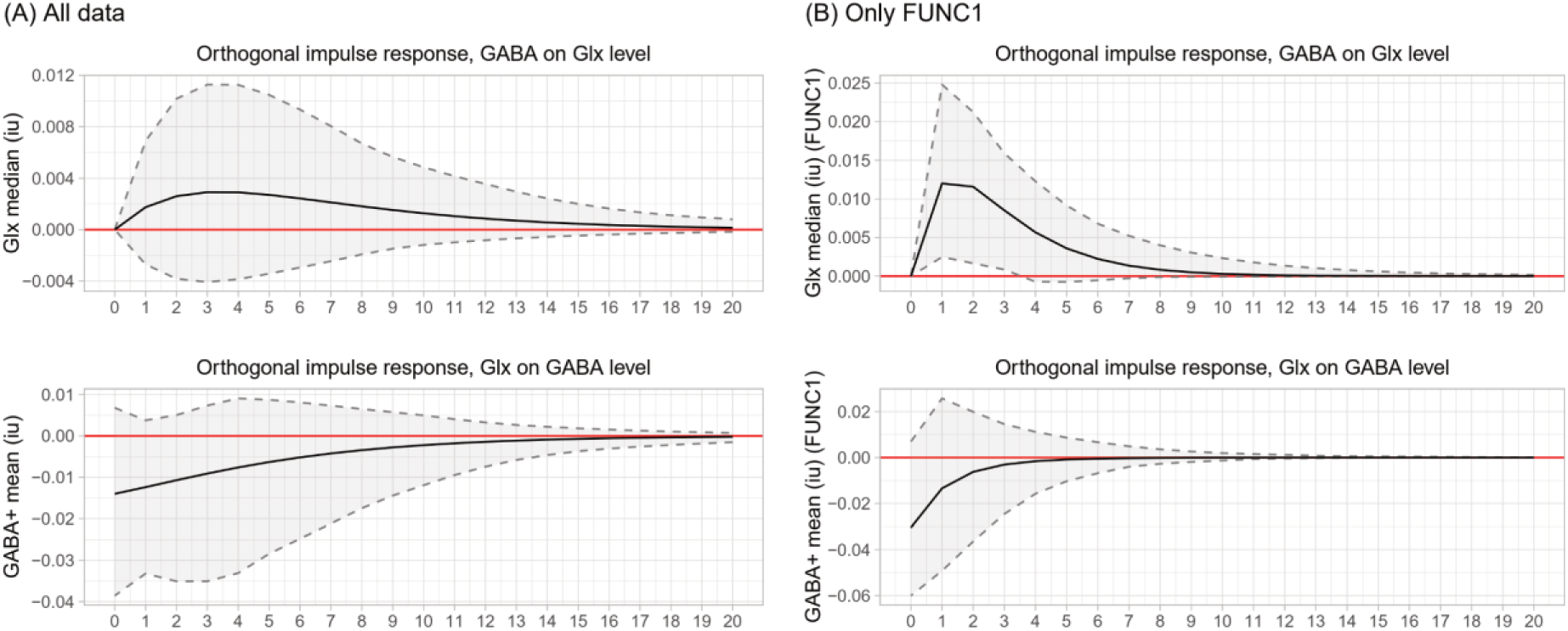
Impulse response function (IRF) of VAR models included data from (A) all conditions and (B) from only FUNC1 of sliding window analysis. The IRF plot shows the impact of a one-unit change of one variable on another variable (i.e., the effect of one unit of GABA change on Glx evolution). A total of 20 lags were given, where one lag represented 1 transient (2s). The shaded region is 95% confidence intervals.

### Correlation analysis suggests GABA+/Glx decouple during stimulation in FUNC1

Overall, Spearman correlation coefficients showed a consistent trend of decoupling between GABA+ (i.u.) and Glx (i.u.) during the first functional block, where the Spearman correlation transitioned from a positive to a negative value in FUNC1. For block analysis, weak negative correlations were observed but were not statistically significant (Fig. 5A). Full correlation details shown are in Supplementary Table 6. When considering sub-block analysis with higher temporal resolution, the data indicated a statistically significant correlation between GABA+ (i.u.) and Glx (i.u.) towards the end of the first stimulation block (FUNC1ON4; R = −0.54, *p* = 0.02) and in the middle of the second stimulation block (FUNC2ON5; R = 0.50, *p* = 0.03) (Fig. 5B). Full correlation details for subblock analysis are available in Supplementary Table 7.

**Figure 5.**
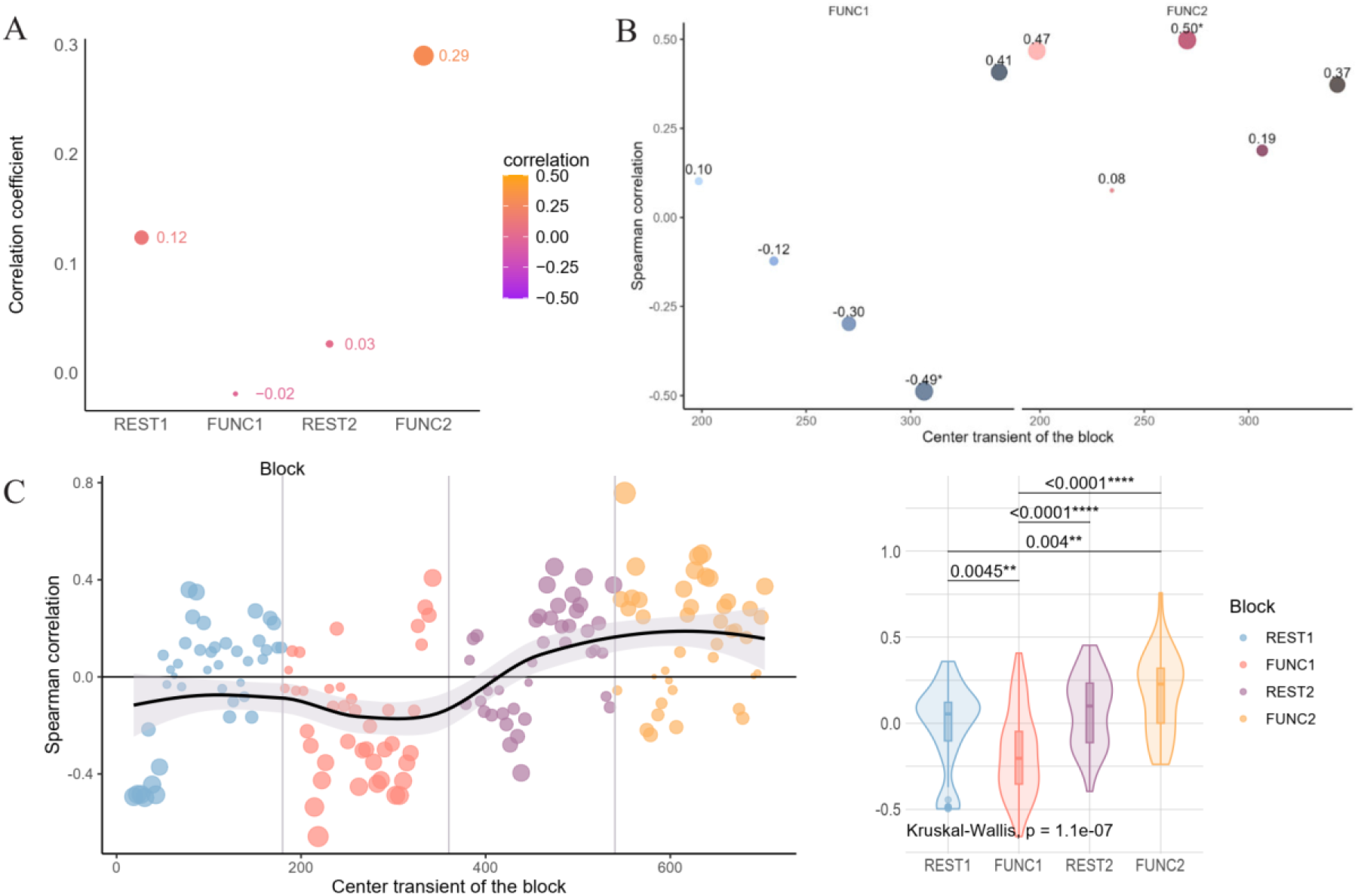
Spearman correlations between GABA+ (i.u.) and Glx (i.u.) for each analysis approach. (A) For each block obtained through block analysis, (B) For each subblock, where * represents Spearman correlation with statistically significant and violin plot of Spearman correlations with the P-value obtained from Wilcoxon signed rank test. (C) Spearman correlation for each timepoint of the sliding window analysis. The black line shows a non-linear fit based on the LOESS fitting method, with the grey ribbon representing the 95% confidence interval. For all plots, the size and number above each dot represent the magnitude of the correlation.

This same trend of GABA+ (i.u.) and Glx (i.u.) decoupling was consistently observed in sliding window analysis data, where correlation coefficients between GABA+ (i.u.) and Glx (i.u.) were mostly negative during FUNC1. A Kruskal-Wallis test showed a significant difference in the Spearman correlation coefficient (*p* =1.1×10^-6^). Post-hoc Wilcoxon signed-rank tests adjusted for multiple comparisons showed statistically significant differences between REST1– FUNC1 (*p* = 0.0044), REST1–FUNC2 (*p* = 0.0044), FUNC1–REST2 (*p* = <0.0001), and FUNC1–FUNC2 (*p* = <0001) (Fig.5C) thereby suggesting constant modulation between GABA+ and Glx throughout the experiments.

We observed similar GABA-Glx decoupling with correlation coefficients calculated using GABA+ and Glx concentrations derived from the fitted peak area (to avoid quantification relative to water) instead of tissue-corrected concentrations (Supplementary Fig. 5–7). While we did not observe a significant correlation toward the end of the first fMRS block for subblock analysis (FUNC2ON5), as seen with tissue-corrected metabolites, the same decoupling trend suggests that the choice of reference did not influence the results.

We also investigated the Glx/GABA+ ratio (Supplementary Fig. 8–10), however, we did not find any statistically significant results for any of the analysis pipelines.

### Metabolite levels at rest negatively correlate with the magnitude of metabolite changes

Overall, the correlations demonstrated statistically significant negative correlations between the levels of GABA & Glx at REST1 and the %change in metabolite levels for both block and subblock analyses (Fig. 6). Moderately negative correlations between metabolite levels at REST1 and metabolites in other blocks were observed for GABA+ (i.u.) (R = −0.44, *p* = 0.00189), and Glx (i.u.) obtained with block analysis (R = −0.38, *p* = 0.00108) (Fig. 6). Subblock analysis showed a weak negative correlation for both GABA+ (i.u.) (R = −0.26, *p* = 0.0028), and Glx (i.u.) (R = −0.31, *p* = 0.00021). These results provide support that baseline metabolite levels correlate with subsequent changes in metabolite levels in response to tactile stimulation.

**Figure 6.**
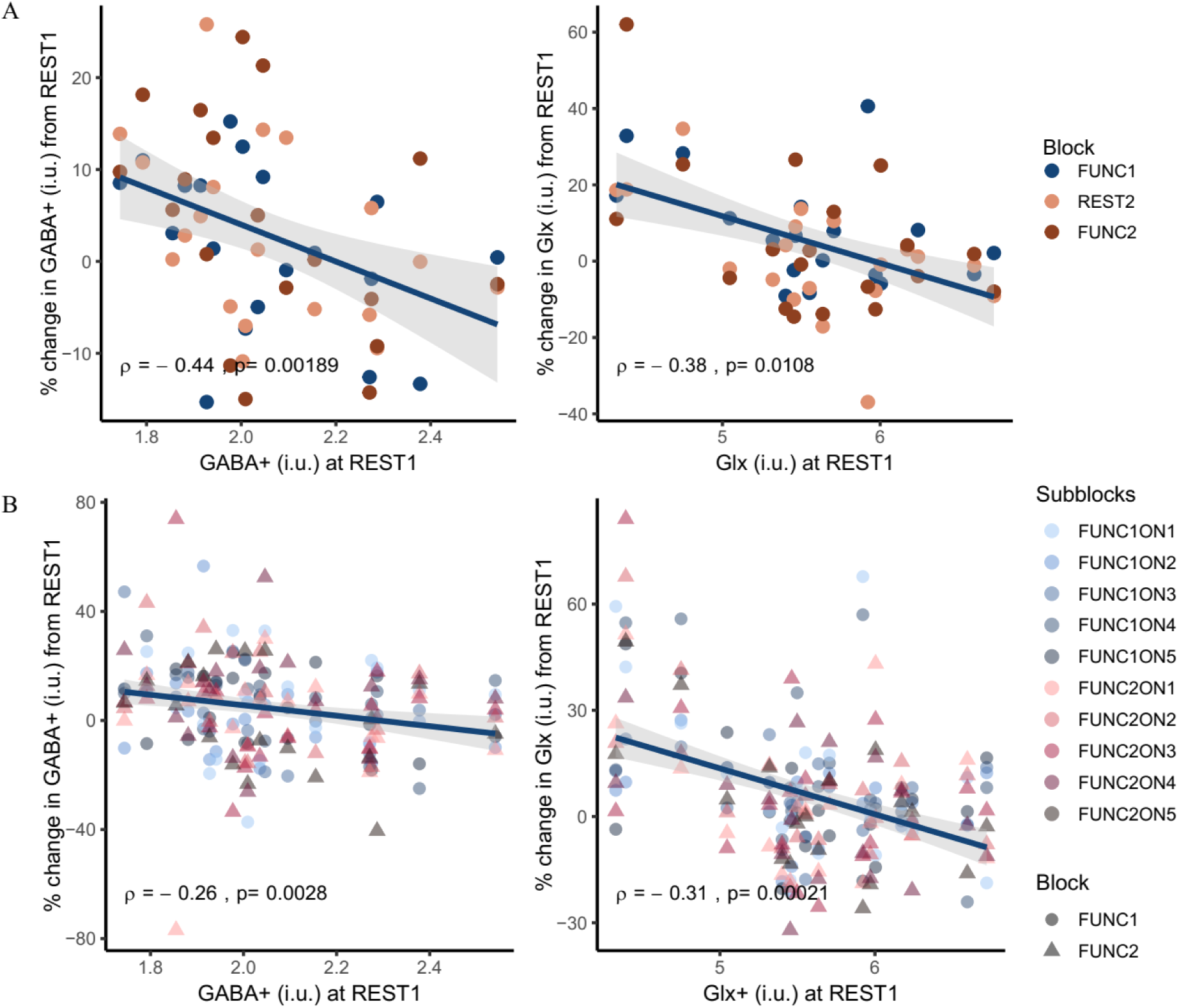
Correlations between the levels of GABA and Glx at REST1 and the %change in metabolite levels for both (A) block and (B) subblock analyses. Spearman correlations of metabolites quantified using block analysis at REST1 compared to other conditions. P-values were corrected for multiple comparisons using the FDR method. Linear lines are plotted purely for visualisation purposes to show directionality.

### GABA+ and Glx correlate with tactile perception

For block analyses, there were significant moderate negative correlations of Glx %change across conditions with our perceptual marker of feedforward inhibition (ρ = −0.46, *p* = 0.003), dynamic detection thresholds (R = −0.42, *p* = 0.003), and amplitude discrimination thresholds (ρ = −0.39, *p* = 0.006). When examining specific blocks, the Glx %change during FUNC1 showed a significant moderate correlation with the dynamic detection threshold (R = −0.42, *p* = 0.003) (Fig. 7). For subblock analysis, weak positive correlations were observed for the %change in GABA+ across conditions with the static detection threshold (ρ = 0.25, *p* = 0.01) and a significant weak positive relationship between Glx %change and tactile adaptation (ρ = 0.3, *p* = 0.01) (Fig. 7). No significant correlations were observed when focusing on a specific subblock.

**Figure 7.**
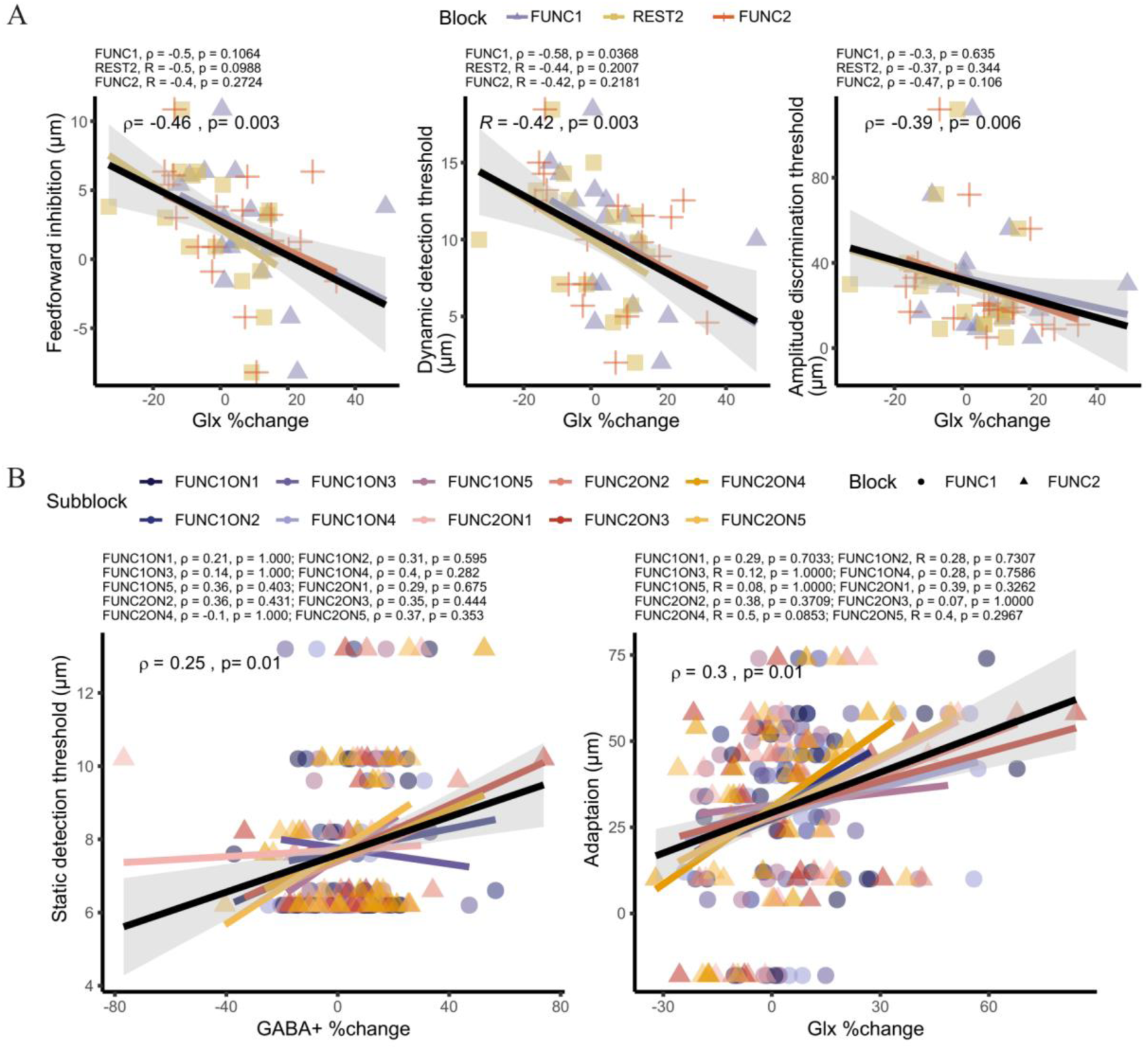
Linear regression between Glx %change and vibrotactile measures. (A) Results for block analysis. (B) Results for subblock analysis. The overall correlations between the variables are displayed on each plot, with correlations between metabolites during specific conditions and vibrotactile measures noted at the top. P-values were corrected for multiple comparisons using the FDR method. Only significant correlations are displayed.

## Discussion

Here we explored the effects of perceptual adaptation through repetitive tactile stimulation on GABA+ and Glx dynamics using functional MRS. Although we did not identify any statistically significant changes in individual metabolites (GABA+ and Glx) at the level of stimulus block, we did find a significant negative correlation *between* GABA+ and Glx in the first stimulus block. Subsequently, multivariate time series analysis of the first stimulus block, demonstrated that the inclusion of the Glx time series enhanced the prediction of the GABA+ time series. Critically, however, while GABA and Glx were positively correlated at baseline (as many other studies at baseline have shown,^68,69^) they decoupled during stimulation. Thus, rather than stimulation leading to gross changes in metabolite concentration, the metabolites instead shift in relation to each other. Interestingly, metabolite levels were also significantly correlated with measurable behavioural outcomes of tactile perception, some of which were in line with our previous findings.^70–72^ In summary, we find a complex temporal interaction between GABA and Glx in response to repetitive tactile stimulation, showing that the change in metabolite levels are both dependent on each other, and that experience-dependent plasticity is reflected in a decoupling of GABA and Glx function.

First, our data reveal a dynamic interaction between GABA+ and Glx under functional conditions in an intercorrelated manner. VAR analysis demonstrated that during the first stimulus block (FUNC1), the inclusion of the Glx time series in the model significantly improved the prediction of the GABA+ time series. In agreement with this notion, IRF plots showed that only for the first stimulation block, the impulse reflecting an increase in GABA+ led to significant increases in Glx levels 6s after GABA+ increases which then returned to baseline. In agreement with many studies, this suggests that homeostatic regulation of GABA and glutamatergic processes is necessary for normal sensory processing.^6,8,73^ This result might also suggests that changes in metabolites occur in a transient manner, with a possible lag following stimulus presentation, as proposed by many fMRS studies.^31,32^ How and why this only occurs during FUNC1 remains unclear. It is possible that the brain is habituating/adapting during the first stimulation block as an iterative process, by which GABA impulses subsequently modulate the release of glutamate to retain homeostatic control. Whereas this mechanism is less implicated either at rest condition, or when, in the case of the second stimulation block perhaps, the brain is already in an adapted state.

We also observed significant negative correlations between metabolite levels at rest and the magnitude of metabolite level changes across conditions. Specifically, the higher the GABA+ or Glx level at baseline, the greater the decrease across both resting and functional conditions. These results agree with previous studies showing a negative relationship between baseline GABA levels and percentage changes in GABA levels after a motor learning task.^74,75^ This also suggests a regulatory homeostatic mechanism for GABA+ and glutamate, where shifts in these metabolites in response to stimulation are tightly controlled and will regress to a homeostatic state. ^74^ Taken together with our VAR approach, we are now able to show the temporal dynamics of this process which appears to be on the order of around 6 seconds.

Beyond this temporal association, correlation analysis revealed a decoupling of GABA+ (i.u.) and Glx (i.u.), where their correlation turned negative during the first fMRS block. These decoupling correlations were significant toward the end of the first fMRS block and were also consistent with the sliding window analysis, which demonstrated significant differences in correlation coefficient values between different stimulation conditions, internally validating this finding with two analytical approaches. Large cohort studies across the lifespan have shown a positive correlation between GABA (i.u.) and Glx (i.u.) at rest.^68,69,76^ Here, we can infer that the observed decoupling or shift to negative correlation values between GABA (i.u.) and Glx (i.u.) in the current study is exclusively driven by the repetitive tactile stimulation applied.

This decoupling potentially suggests a shift in excitation-inhibition balance in response to tactile stimulation, where the overall activity of neurons facilitates plasticity by modulating different temporal rates of change in excitation and inhibition, which results in an anticorrelation during the first functional block. A number of studies suggest a role of GABAergic disinhibition as one of the necessary factors facilitating learning in many domains.^38,77–80^ Upregulation of glutamatergic synaptic activity and glutamate levels appear correlated with learning efficiency and tactile activation.^37,81,82^ Another possible explanation is the temporary demand for GABA, which may lead to a temporary reduction in Glx levels due to the conversion of glutamate to GABA, explaining the negative correlation observed.

The return to positive correlation values in the second rest block (REST2) suggests tight regulation of excitation-inhibition homeostasis, favouring a positive correlation between GABA+ (i.u.) and Glx (i.u.). This shift, beginning immediately after the significant negative correlation observed during the first functional block, suggests that the process may occur on a timescale of minutes, as shown by the subblock analysis with an effective temporal resolution of 72 s (36 transients). The absence of a negative correlation in the second stimulus block (FUNC2), which we hypothesise results from repetitive tactile stimulation, suggests potential brain adaptation in response to the stimulus and is consistent with the VAR results.

We should note that a positive relationship between GABA+ and Glx at rest was also observed in parietal, prefrontal, and occipital cortex in prior work.^68,69,76^ However, another study failed to find a positive correlation between GABA+ and Glx in the motor and visual cortex,^83^ suggesting this may be region specific, although the underlying mechanisms remain unclear. To further confirm the effect of repetitive tactile stimulation on GABA+/Glx correlations, or GABA+/Glx correlations in general, future investigations on this topic are warranted.

Disturbances in either glutamatergic or GABAergic systems have been suggested to underlie sensory processing differences in many neurodevelopmental conditions. For example, higher glutamate + glutamine was associated with higher reactivity in autistic children^84^ and adults^85^, yet other studies have shown that reduced GABA drives sensory differences.^34,86–88^ In our study, the correlations between GABA+ (i.u.) and Glx (i.u.) levels and various vibrotactile measures indicate a complex relationship between metabolite levels and sensory processing. While prior work has shown such relationships, they tended to look at metabolites at rest rather than “under stress” by stimulation.

Here, we found a negative correlation between the overall Glx % change and dynamic detection threshold and amplitude discrimination threshold. Specifically, the greater the increase in Glx across conditions, the lower the dynamic detection threshold and amplitude discrimination threshold, suggesting that increased excitability after stimulation (e.g. more responsiveness of the system) is associated with an increased ability to detect or discriminate between subtle vibrations, essentially increasing the gain of the system. When examining finer temporal resolution with sub-block analysis, we found a weak positive correlation between Glx %change and the difference between amplitude discrimination thresholds with and without adaptation, which serves as a proxy for adaptation processes.^42^ Specifically, the higher the Glx increase, the higher the adaptation. Given that Glx is associated with cortical excitability,^89^ this result is in agreement with previous studies linking higher cortical excitability with motor learning and adaptation.^81,90,91^

While we focus on Glx, these results align with the notion that glutamate is necessary for sensory processing, and alterations in glutamate levels have been linked to sensory reactivity in e.g. autism.^84,92^ One MRS-EEG study indicated that an increase in glutamate levels is associated with lower prediction errors to deviant stimuli,^93^ and subsequent research showed that higher Glx is associated with stronger connectivity during unpredictable stimulation.^94^ Our study also revealed that a stronger reduction in Glx levels corresponds to a higher feedforward inhibition measure, reflecting more inhibition (the threshold is expected to increase after pre-pulse stimulation when there is more inhibition). This is consistent with previous studies in children.^84^ These findings highlight the importance of glutamate in modulating neuronal excitation and preventing overexcitation.^95^ We also found that GABA+ % change across sub-blocks was weakly associated with static detection thresholds. The static detection threshold is a well-known measure for GABAergic processes, sensory gating in particular, and normal brain function,^42,85,87^ suggesting that our fMRS-measured GABA+ reflects its role in sensory processing. Furthermore, % change may reflect the ability of the system to exert inhibition more than baseline measures of GABA. Exactly how GABA might drive these perceptual results remains uncertain and task-based tactile fMRS studies are needed but suffer from low SNR.

In contrast with our hypothesis and previous studies that observed changes in individual metabolite levels in response to repetitive tactile stimulation,^10,37,38,79^ we did not find any gross significant changes in metabolite levels during stimulation blocks. This lack of significant differences was consistent across all analysis pipelines used in the current study. Beyond these signals potentially being averaged out (see^32^), one potential explanation is the stimulus frequency used. It has been demonstrated that neuronal plasticity is influenced by both stimulation frequency and intrinsic cortical network oscillations. Different stimulus frequencies—high (>10 Hz) and low—lead to long-term potentiation (LTP) and long-term depression (LTD), respectively.^21,24,96,97^

This notion of experience-dependent plasticity is supported by studies showing that applying digit co-stimulation at the somatosensory cortex’s resonance frequency of 23 Hz leads to perceptual improvement.^98^ Conversely, below-resonance stimulation at 7 Hz and above-resonance stimulation at 39 Hz result in deteriorated tactile task performance and slower reaction times.^96^ Subsequent fMRS studies have revealed that GABA release during repetitive tactile stimulation is frequency-dependent, with perceptual learning and GABA level reduction occurring exclusively at above-resonance frequencies (39 Hz).^98^ The same study also demonstrated that initial GABA levels predicted the extent of perceptual learning.

Our current experimental design employed repetitive stimulation at a frequency of 3 Hz, as it has been reported that low-frequency stimulation could also drive LTP in a longer time course through the same cellular mechanisms.^24,99,100^ However, the lack of metabolite changes to repetitive tactile stimulation in the current study could suggest that the changes induced by the current stimulus frequency are more subtle compared to other studies that use higher frequencies to induce plasticity. Another possibility is that the plasticity induced by this current stimulus operates mainly through the N-methyl-D-aspartate (NMDA) receptors which act as an activity-dependent coincidence detector.^101–104^ It is well-documented that dynamic changes in NMDA expression allow for the modification of synaptic strength and induction of LTP and LDP.^104–107^ However, this change in NMDA expression does not directly involve a change in glutamate concentration and has been suggested to not be detectable by MRS measurement.^108^

Finally, our study is not without limitations. These include the small number of participants and consequently low effect size, particularly for analyses with a low signal-to-noise ratio (SNR) such as sub-block and sliding window analysis. However, MRS data quality assessment was carefully carried out to ensure the reliability of the included data points and our multi-method approach to analysis internally validates our VAR and decoupling of GABA and Glx findings. Another limitation is that we did not measure the psychophysical metrics post-fMRS experiment to confirm adaptation to repetitive tactile stimulation. This decision was partly due to the desire to avoid participant fatigue from long experimental procedures, which could influence the results, and the presence of fMRI tasks after the fMRS (included for a different study) which likely would have reduced the impact on repetitive stimulation, which is short-lived. Future studies with shorter fMRS or psychophysics tasks would be of interest to further investigate sensory processing through tactile adaptation, directly linking changes in metabolites to *changes* in perception.

## Conclusion

Our study suggested that decoupling between GABA and Glx is induced exclusively by tactile stimuli, alongside the predictive relationship and impulse response findings between these two neurotransmitters. Our results not only show the feasibility of fMRS technique for studying brain responses during tactile processing, but also offer novel insights into the functional coupling and the temporal relationship between GABA and glutamate during stimulation. Our results suggest that, instead of focusing separately on GABA+ for inhibition and glutamate for excitation, future studies would benefit from exploring how these neurotransmitters dynamically coordinate, to gain a better understanding of their role in tactile processing.

## Data availability

Data is available upon reasonable request.

## Acknowledgements

We would like to thank all the participants who took part in this study.

## Supplementary material

Supplementary material is available at *Brain* online.

## Abbreviations

ADHD: attention deficit hyperactivity disorder
E/I balance: Excitation inhibition balance
fMRS: functional Magnetic Resonance Spectroscopy
Glx: Glutamine + Glutamate
i.u.: Institutional units
MRS: Magnetic Resonance Spectroscopy

