## Supplementary Figure for "Decoupling of GABA and Glutamine-Glutamate Dynamics and their role in tactile perception: An fMRS Study"

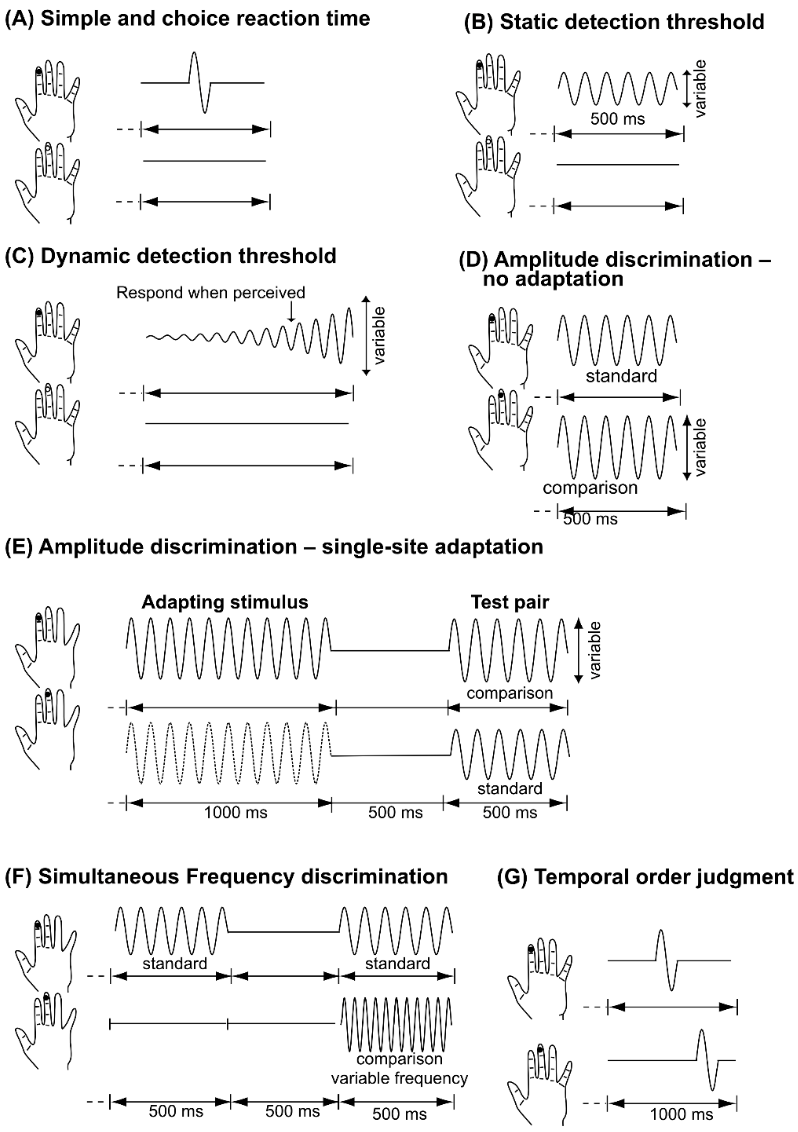


Supplementary Figure.1 (A) Simple and Choice reaction time, (B) Static and (C) Dynamic detection threshold, (D) Amplitude discrimination without adaptation and (E) with single-site adaptation, (F) Simultaneous frequency discrimination, and (G)Temporal Order Judgment. Adapted from a publicly available figure on https://osf.io/2ygpc.

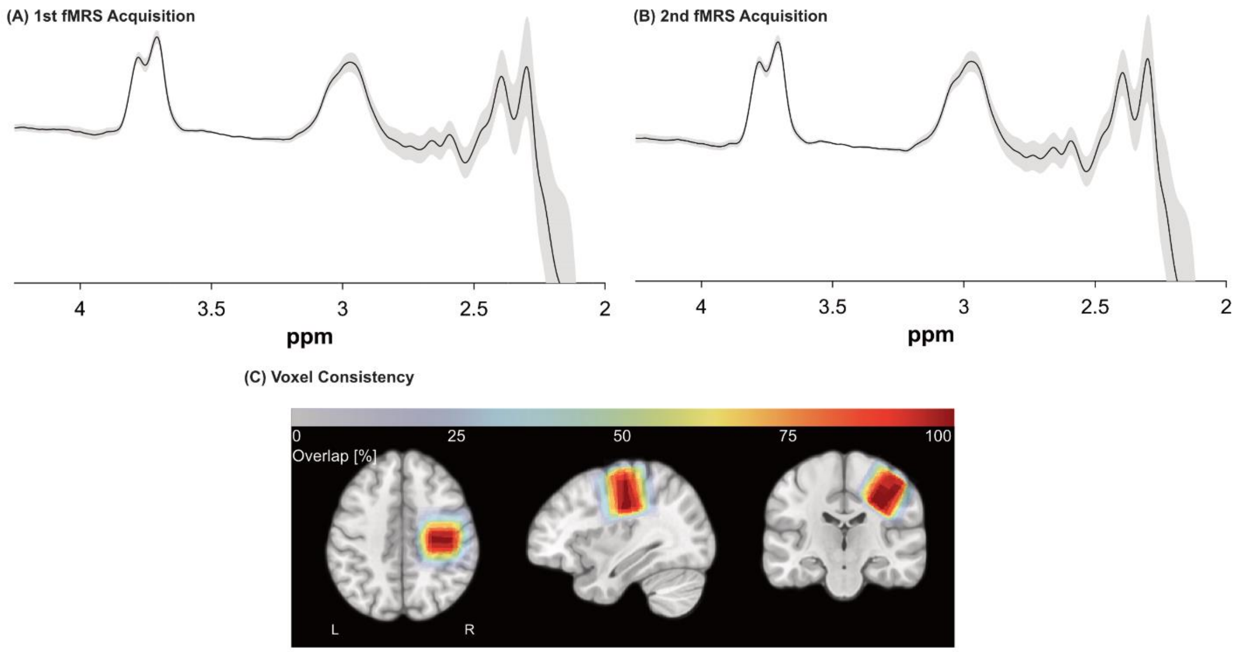


Supplementary Figure.2 Averaged MEGA-PRESS difference spectra from (A) 1st fMRS acquisition and (B) 2nd fMRS acquisition. The grey shading on (A) and (B) represents 95% CI. (C) Voxel positioning consistency across all participants.

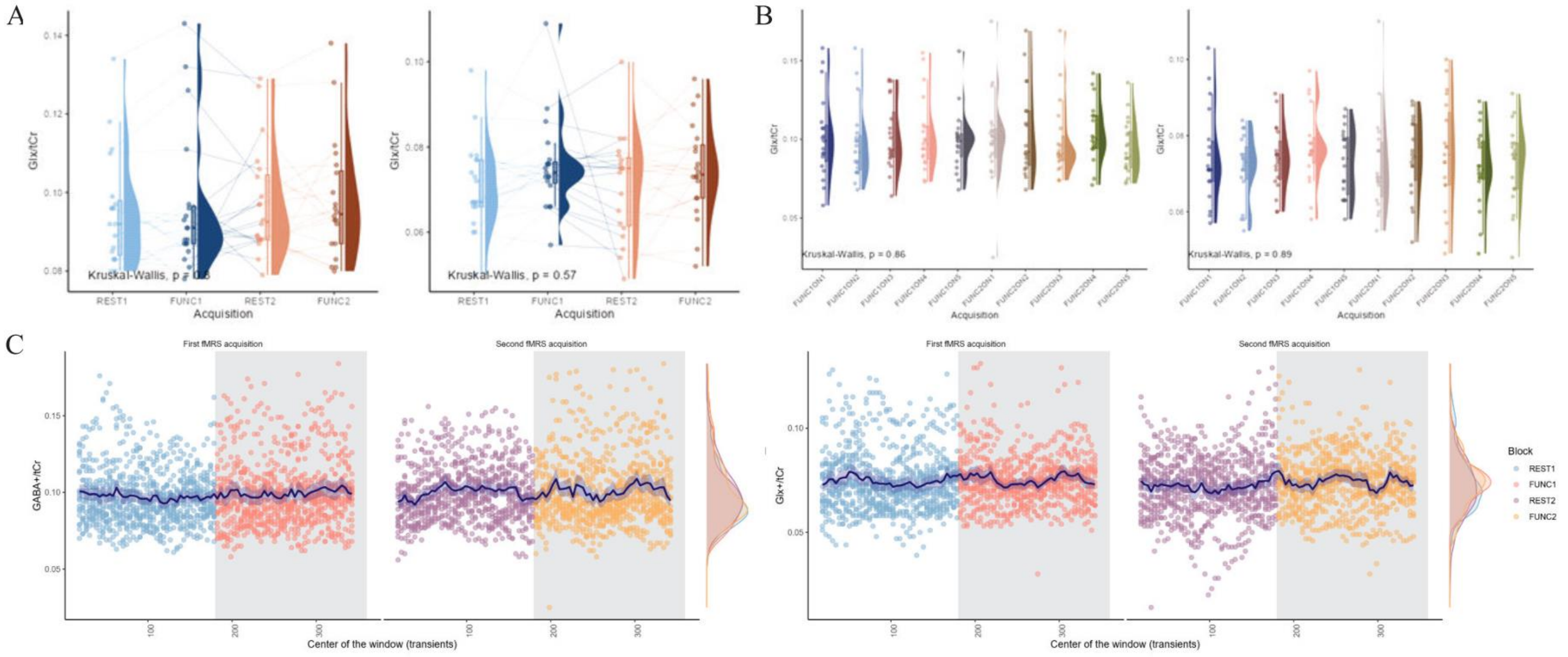


Supplementary Figure.3 Raincloud plot of (A) block and (B) subblock analysis and (D) sliding window analysis results. The lines connect data points within the same participant.

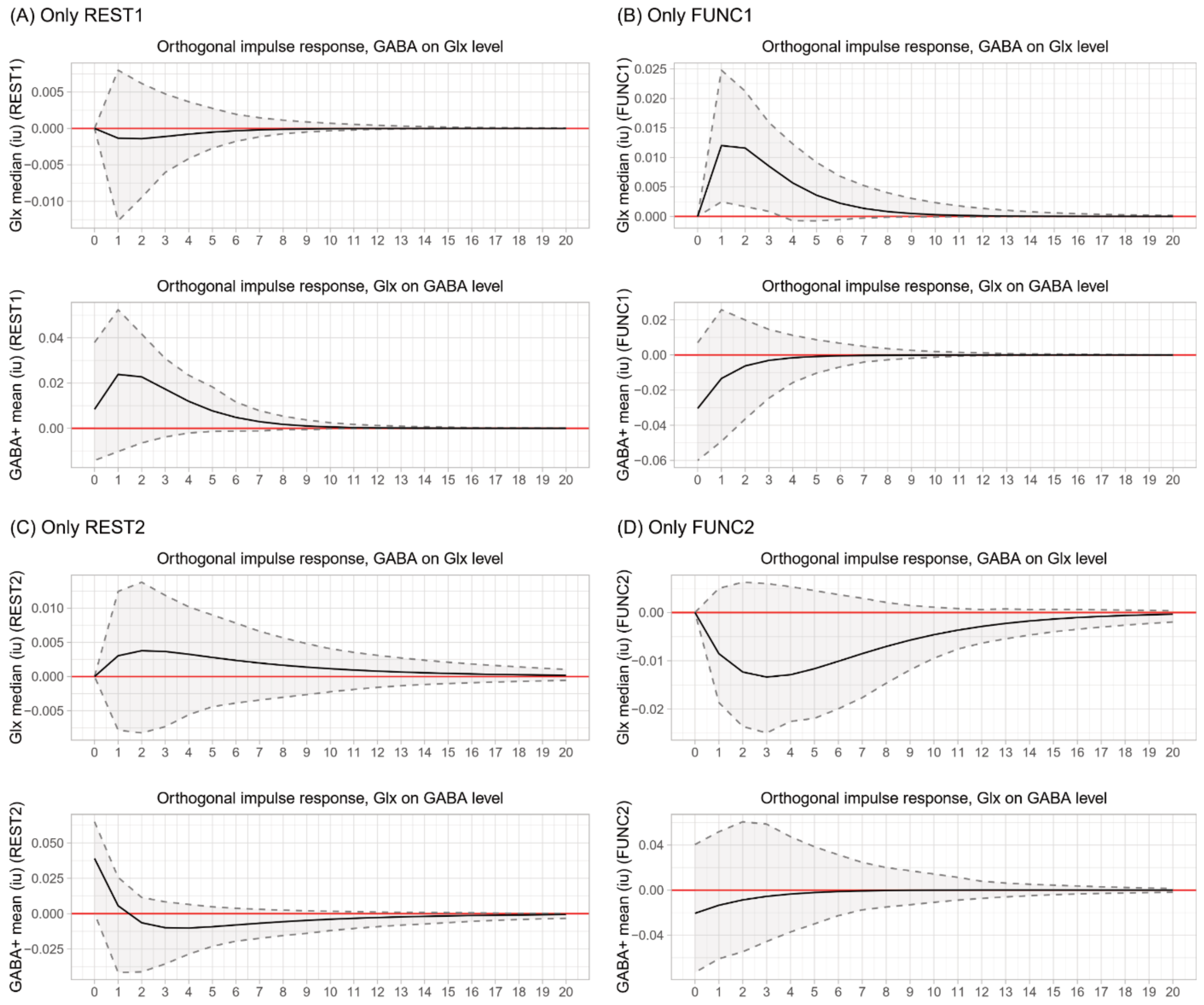


Supplementary Figure.4 Impulse response function (IRF) of VAR models included data from (A) only REST1, (B) only FUNC1, (C) only REST2, (D) only FUNC2 of sliding window analysis. A total of 20 lags were given, where one lag represented one transient (2 s). The IRF plot shows the impact of a one-unit change of one variable on another variable (i.e., the effect of one-unit of GABA change on Glx evolution). The shaded region is 95% confidence intervals.

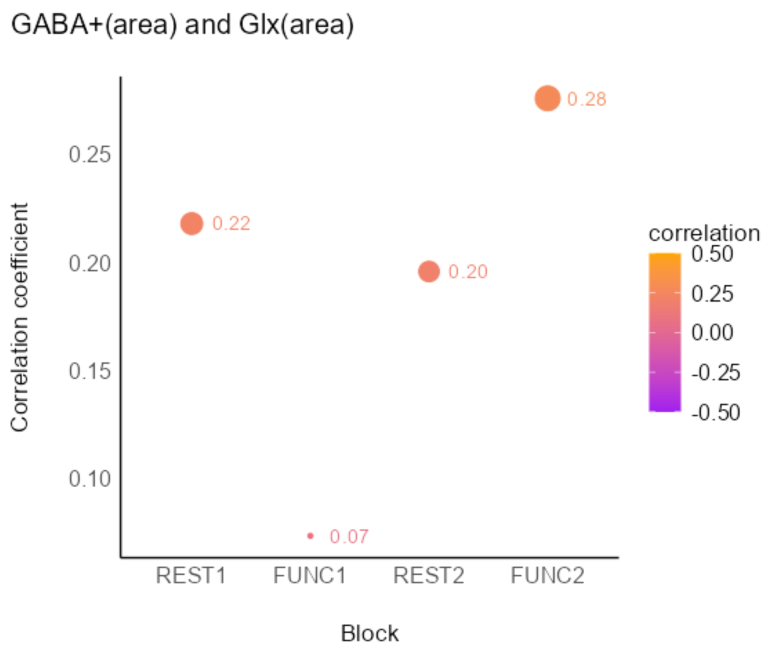


Supplementary Figure.5 Spearman correlations between GABA+ and Glx concentrations obtained from fitted peak area for each block obtained through block analysis. The size of each dot represents the magnitude of the correlation.

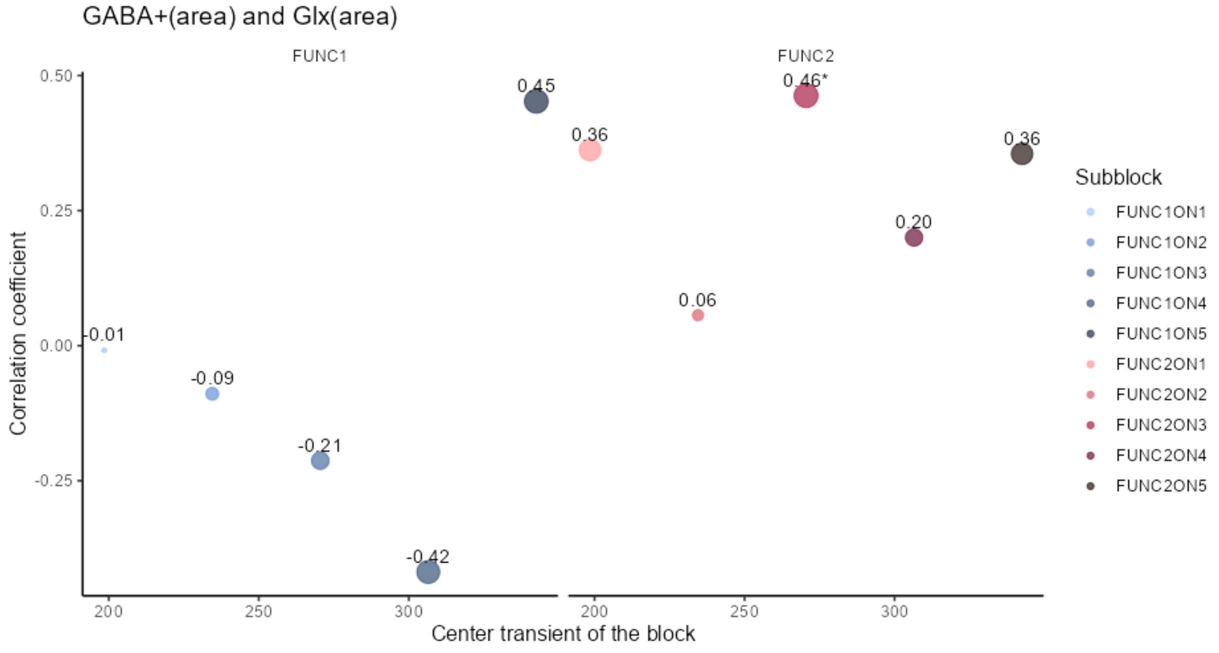


Supplementary Figure.6 Spearman correlations between GABA+ and Glx concentrations obtained from fitted peak area for each subblock. The size and number above each dot represent the magnitude of the correlation, where * represents statistically significant Spearman correlation at p<0.05

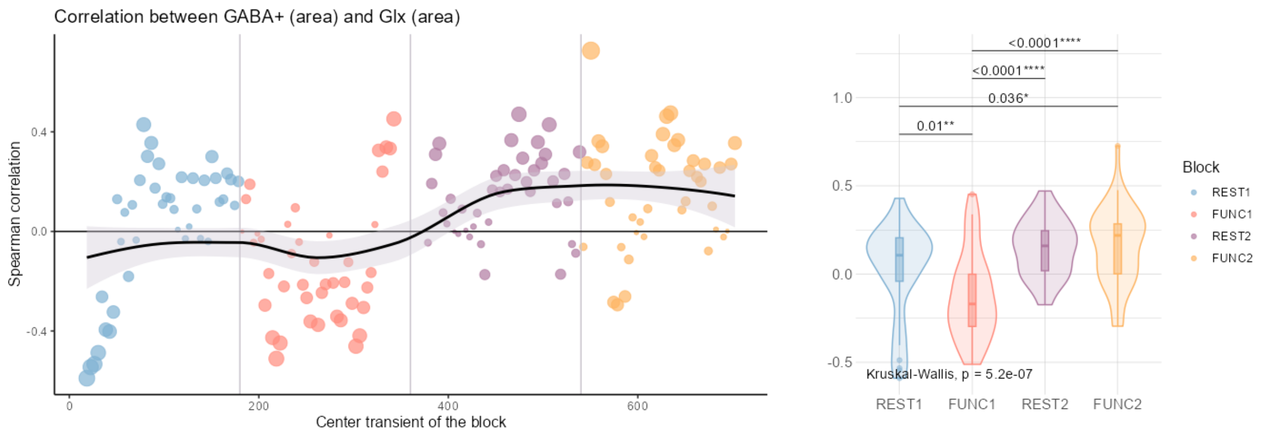


Supplementary Figure.7 (Left) Spearman correlations between GABA+ and Glx concentrations obtained from fitted peak area for each timepoint in sliding window analysis. Non-linear fit based on the LOESS fitting method, with the grey ribbon representing the 95% confidence interval. The size of each dot represents the magnitude of the correlation. (Right) Violin plot of Spearman correlations. The p-value is obtained from the Kruskal-Wallis test, with pair-wise post-hoc test p-values obtained from the Wilcoxon signed-rank test, corrected for multiple comparisons using the FDR method.

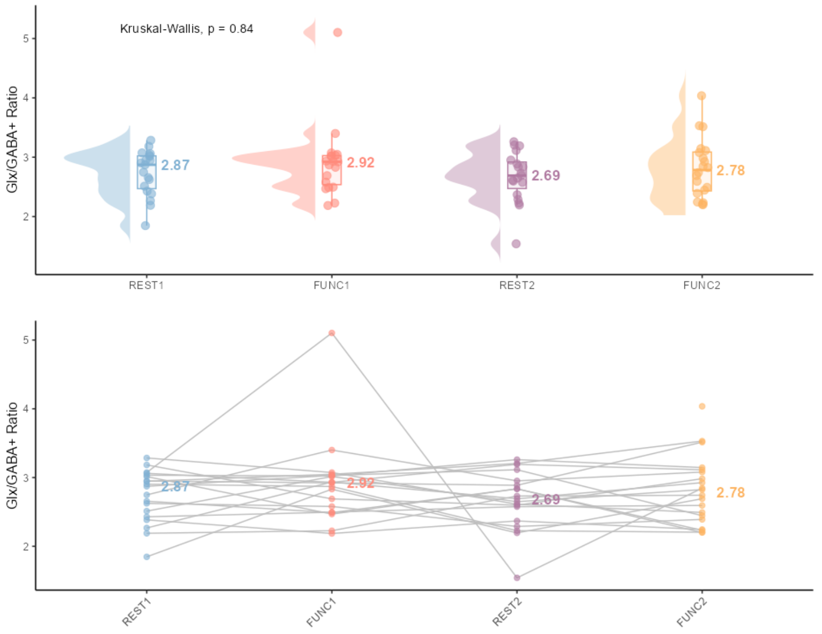


*Supplementary* Figure.8 *Raincloud plot of Glx/GABA+ ratio from block analysis results. The lines connect data points within the same participant.*

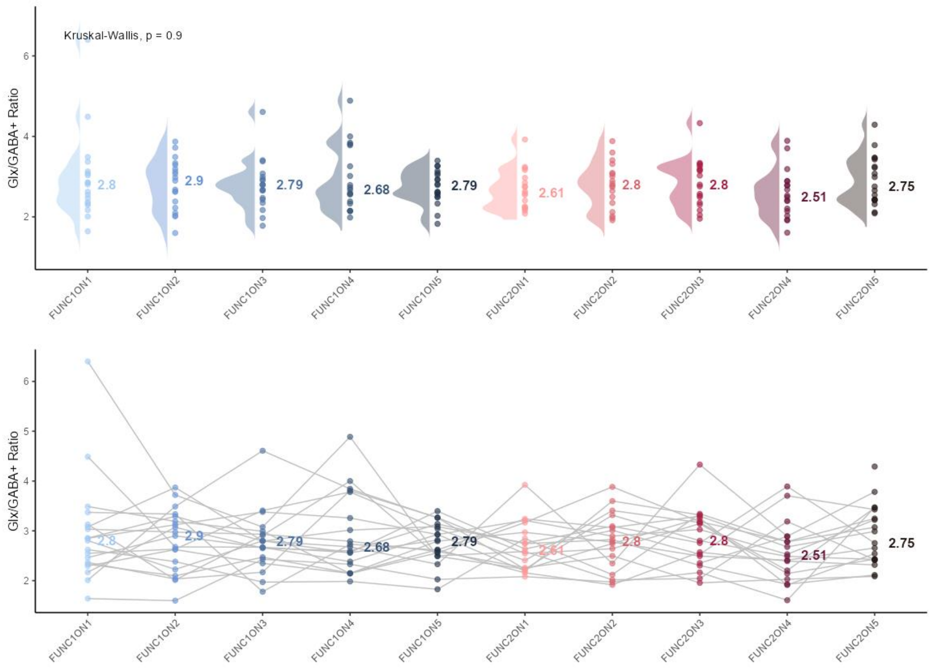


*Supplementary* Figure.9 *Raincloud plot of Glx/GABA+ ratio from subblock analysis results. The lines connect data points within the same participant.*

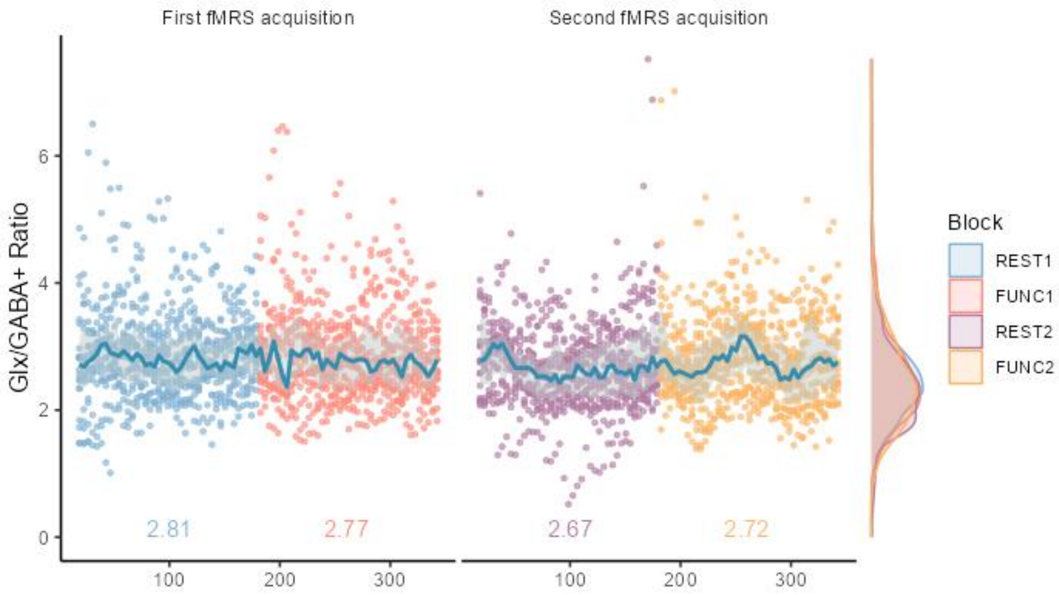


Supplementary Figure.10 Dotplot of sliding window analysis. The x-axis represents the centre of the window for metabolite quantification for each block, where each acquisition includes a total of 320 transients. Blueline/Ribbon = median+ lower and upper quartiles.

**Supplementary Table 1. MRSinMRS checklist**

| Site (Name or Number) | |
|---|---|
| **1. Hardware** | |
| **a. Field strength [T]** | 3 T |
| **b. Manufacturer** | GE |
| **c. Model (software version if available)** | Signa Premier (software version MR30.1_R01_2322.c) |
| **d. RF coils: nuclei (transmit/ receive), number of channels, type, body part** | 48-channel head coil receiver |
| **e. Additional hardware** | |
| **2. Acquisition** | |
| **a. Pulse sequence** | MEGA-PRESS |
| **b. Volume of Interest (VOI) locations** | Somatosensory cortex |
| **c. Nominal VOI size [$cm^3$, $mm^3$]** | 3x3x3 $cm^3$ |
| **d. Repetition Time (TR), Echo Time (TE) [ms, s]** | TE/TR = 68/2000 ms |
| **e. Total number of Excitations or acquisitions per spectrum** **In time series for kinetic studies** **Number of Averaged spectra (NA) per time-point** | **Total number of spectra acquire**: 2 spectra acquired per participant, 180 transients each<br>**Number of average transients per time point**:<br>Block analysis: 180 transients; Total 4 timepoints from 2 spectra acquired<br>Subblock analysis: 36 transients; Total 10 timepoints from 2 spectra acquired |

| | |
|---|---|
| **Averaging method (e.g. block-wise or moving average)** | Sliding window analysis: 40 transients for each timepoint, sliding step of 4 transients; Total 164 timepoints from 2 spectra acquired |
| **Total number of spectra (acquired / in time-series)** | All spectral analyses were carried out with Gannet version 3.3.2 using default settings except for trimming of transients was used based prior to preprocessing step. |
| **f. Additional sequence parameters (spectral width in Hz, number of spectral points, frequency offsets)** | Data points = 2048, Spectral width = 2000 Hz, frequency offset = -2 ppm |
| **g. Water Suppression Method** | CHESS |
| **h. Shimming Method, reference peak, and thresholds for “acceptance of shim” chosen** | Automatic shimming, Acceptable threshold: < 10 Hz |
| **i. Triggering or motion correction method (respiratory, peripheral, cardiac triggering, incl. device used and delays)** | - |
| **3. Data analysis methods and outputs** | |
| **a. Analysis software** | Gannet version 3.3.2 |
| **b. Processing steps deviating from quoted reference or product** | Modified *Gannetload* function in Gannet for cutting up transients for analysis. |

**c. Output measure (e.g. absolute concentration, institutional units, ratio)Processing steps deviating from quoted reference or product**

- In ratio to creatine
- In institutional units (i.u.) in ratio to unsuppressed water signal within same voxel and corrected for tissue fraction, considering intrinsic metabolites differences in gray matter and white matter (Harris et al., 2015).

**d. Quantification references and assumptions, fitting model assumptions**

**4. Data Quality**

**a. Reported variables (SNR, Linewidth (with reference peaks))**

| Quality metrics | SNR | FWHM (Hz) | Glx Fit Error% | GABA Fit Error% |
|---|---|---|---|---|
| Block | 293.08 ± 80.2 | 9.37 ± 1.63[a] | 3.23 ± 1.21 | 3.63 ± 1.39 |
| Subblock | 152.36 ± 31.61 | 9.38 ± 1.02 | 5.41 ± 2.11 | 5.95 ± 3.04 |
| Sliding window | 151.06 ± 39.13 | 9.38 ± 1.05 | 5.39 ± 2.06 | 5.88 ± 2.21 |

**b. Data exclusion criteria**

Excluded the quantified metabolites if the FWHM of NAA was > 20 Hz or the % fit error of that metabolite was more than 65%.

Extreme outliers were then identified and removed using a boxplot method, where values above the third quartile (Q3) + 3×interquartile range (IQR) or below the first quartile (Q1) - 3×IQR were considered extreme outliers.

**c. Quality measures of postprocessing Model fitting (e.g. CRLB, goodness of fit, SD of residual)**

Model fitting error

**d. Sample Spectrum**

In figure 6.3

**Supplementary Table 2. Descriptive statistics of participants demographics and questionnaire scores.**

| Demographics | N | Mean ± SD |
|---|---|---|
| **Age** | 20 | 25.75 ± 5.8 years old |
| **Sex F (M)** | 12 (8) | - |
| **Handedness R (L)** | 20 (0) | - |
| **AQ-10** | 20 | 2.55 ± 1.6 |
| **GSQ score** | 20 | 41.75 ± 18.3 |

**Supplementary Table 3. MRS quality metrics**

| Quality metrics | SNR | FWHM (Hz) | Glx Fit Error% | GABA Fit Error% |
|---|---|---|---|---|
| **Block** | 293.08 ± 80.20 | 9.37 ± 1.63[a] | 3.23 ± 1.21 | 3.63 ± 1.39 |
| **Subblock** | 152.36 ± 31.61 | 9.38 ± 1.02 | 5.41 ± 2.11 | 5.95 ± 3.04 |
| **Sliding window** | 151.06 ± 39.13 | 9.38 ± 1.05 | 5.39 ± 2.06 | 5.88 ± 2.21 |

FWHM: Full width of half maximum, SNR: Signal to Noise ratio, SNR and FWHM are from NAA peak. All data reported as median ± IQR except for [a] denotes data reported as mean ± SD.

### Supplementary Table 4. Posterior distribution from Bayesian linear-mixed model for each model from sliding window analyses

| Parameter | Median | 95% CI | pd | Significance[a] | Large[b] | Rhat | ESS |
|---|---|---|---|---|---|---|---|
| **Glx (i.u.)** | | | | > \|0.05\| | > \|0.30\| | | |
| **(Intercept)** | | | | | | | |
| **FUNC1** | 0.020 | [-0.15, 0.18] | 58.53% | 34.24% | 0.02% | 1 | 9111 |
| **REST2** | -0.070 | [-0.26, 0.12] | 75.76% | 57.45% | 0.91% | 1 | 7361 |
| **FUNC2** | $-3.810\times10^{-04}$ | [-0.21, 0.21] | 50.12% | 50.12% | 0.38% | 1 | 7871 |
| **GABA+(i.u.)** | | | | > \|0.02\| | > \|0.11\| | | |
| **(Intercept)** | | | | | | | |
| **FUNC1** | 0.006 | [-0.05, 0.07] | 58.23% | 35.79% | 0.05% | 1 | 7096 |
| **REST2** | 0.002 | [-0.07, 0.07] | 51.54% | 32.25% | 0.12% | 1 | 7065 |
| **FUNC2** | 0.030 | [-0.05, 0.10] | 75.49% | 58.20% | 1.61% | 1 | 7544 |

[a,b] Based on SEXIT framework, Threshold for Significant (a) and large (b) effect is 0.05*$SD_y$ and 0.3*$SD_y$ where $SD_y$ is the standard deviation of the outcome parameter (Makowski et al., 2019a).

### Supplementary Table 5. Results from Granger causality tests

| Data | Null-Hypothesis | F-test | P-value | Decision |
|---|---|---|---|---|
| **From all conditions** | GABA+=/>Glx | 0.124 | 0.725 | Do not reject |
| | Glx =/> GABA+ | 1.525 | 0.217 | Do not reject |
| **Only REST1** | GABA+=/>Glx | 1.882 | 0.174 | Do not reject |
| | Glx =/> GABA+ | 0.100 | 0.753 | Do not reject |
| **Only FUNC1** | GABA+=/>Glx | 0.034 | 0.854 | Do not reject |
| | Glx =/> GABA+ | 4.152 | 0.045 | Reject |
| **Only REST2** | GABA+=/>Glx | 0.745 | 0.391 | Do not reject |
| | Glx =/> GABA+ | 0.320 | 0.573 | Do not reject |
| **Only FUNC2** | GABA+=/>Glx | 0.013 | 0.910 | Do not reject |
| | Glx =/> GABA+ | 1.379 | 0.244 | Do not reject |

**Supplementary Table 6. Correlation tests between GABA+(i.u.) and Glx(i.u.) obtained from block analysis**

| **Block** | **Method** | **Correlation coefficient** | ***p*** |
|---|---|---|---|
| **REST1** | Pearson | 0.12 | 0.61 |
| **FUNC1** | Spearman | -0.02 | 0.94 |
| **REST2** | Pearson | 0.03 | 0.92 |
| **FUNC2** | Pearson | 0.29 | 0.21 |

**Supplementary Table 7. Correlation tests between GABA+(i.u.) and Glx(i.u.) obtained from subblock analysis**

| **Subblock** | **Method** | **Correlation coefficient** | ***p*** |
|---|---|---|---|
| **FUNC1ON1** | Spearman | 0.10 | 0.67 |
| **FUNC1ON2** | Pearson | -0.12 | 0.62 |
| **FUNC1ON3** | Pearson | -0.30 | 0.21 |
| **FUNC1ON4** | Pearson | -0.49 | 0.03* |
| **FUNC1ON5** | Pearson | 0.41 | 0.08 |
| **FUNC2ON1** | Spearman | 0.47 | 0.05 |
| **FUNC2ON2** | Pearson | 0.08 | 0.77 |
| **FUNC2ON3** | Pearson | 0.50 | 0.03* |
| **FUNC2ON4** | Pearson | 0.19 | 0.49 |
| **FUNC2ON5** | Pearson | 0.37 | 0.12 |

*$p$<0.05
